# Lunulite bryozoan biogeography – a convergent global success with a distinct Western Australian twist

**DOI:** 10.1101/2023.04.30.538880

**Authors:** Eckart Håkansson, Aaron O’Dea, Antonietta Rosso

## Abstract

Lunulites are a polyphyletic group of marine bryozoans that have been a conspicuous element of marine shelf faunas since the Late Cretaceous into the present day. They are easily recognizable by their domed colony form and free-living mode of life on sea floor sediments. Here we explore the waxing and waning of the major lunulitiform groups and their unique morphology and mode of life from the Cretaceous to the present day. Because relatively few and simple modifications are needed to transition from an encrusting form into this highly specialized lifestyle, shared colonial features are rampant and we find examples of both convergent and iterative evolution across several unrelated clades, although detailed phylogenetic relationship remain largely unresolved. The early chapter of the ‘lunulite story’ is focused on the Late Cretaceous European Chalk Sea, which appears to have been a crucible for the evolution of ‘lunulites’. At least six, and likely more, cheilostome groups independently evolved the free-living mode of life in this tropical shelf area. To what extent any of these free-living clades gave rise to post-Cretaceous groups remains unclear. The Cenozoic chapter is more complex, comprising at least three independently evolved major clades, two of which are extant; (1) the Lunulitidae s. str., a North American/European cluster, comprising the classic *Lunulites*, which became extinct in the late Neogene, (2) the Cupuladriidae, which reached circum-global tropical and sub-tropical distribution in the Miocene, and (3) the ‘Austral lunulite’ cluster, which is almost certainly polyphyletic and through most of its history confined to Australia and New Zealand, bar a comparatively brief colonization of the southern part of South America, with the earliest representatives from NW Western Australia.

## INTRODUCTION

The Bryozoa is a diverse, highly successful Phylum of suspension-feeding, predominantly benthic marine invertebrates with an excellent fossil record dating back to the Early Ordovician, and with a very sporadic, albeit disputed record dating further back, to the beginning of the Cambrian (Zhang et al. 2021, but see also Yang et al 2023). All members of the phylum (with one, possibly derived exception) form colonies by repetitive asexual budding of individual zooids, allowing polymorphic diversification in both shape and function of zooids. Sexual reproduction involves a freely moving larval stage with a duration varying from a few hours to several months (e.g., Driscoll et al. 1971; Cook & Chimonides 1983; Taylor 1988, Winston 1988). The overwhelming majority of bryozoans require a substrate for the larva to successfully settle and metamorphose into the ancestrula or ancestrular complex, thus initiating the new colony. Most bryozoans select stable positions that provide sufficient space to enable the proper development of the colony (Winston 1988; Håkansson & Thomsen 2001). In the case of the free-living bryozoans, however, the lecithotrophic larvae mostly settle on sand-sized grains, typically less than 2 mm in size; far too small to provide stability or space for the adult colony. By outgrowing or integrating the small substrate and adopting heteromorphic zooids called vibracula that possess long, bristle like setae (Fig. 1), the bryozoan colony becomes free-living. The adoption of this free-living mode of life has allowed several groups of cheilostome bryozoans to colonize the vast expanses of fine-grained shelf sediments across the Globe with tremendous success, to the point that in some instances, lunulite bryozoans are the most abundant epifaunal element of benthic soft-sediment communities (Fig. 2) with 2,000 to 3,000 colonies/m^2^ reported by Marcus & Marcus (1962) and an exceptional number of more than 15,000 live colonies/m^2^ reported by Cadée (1975).

**Fig. 1.**
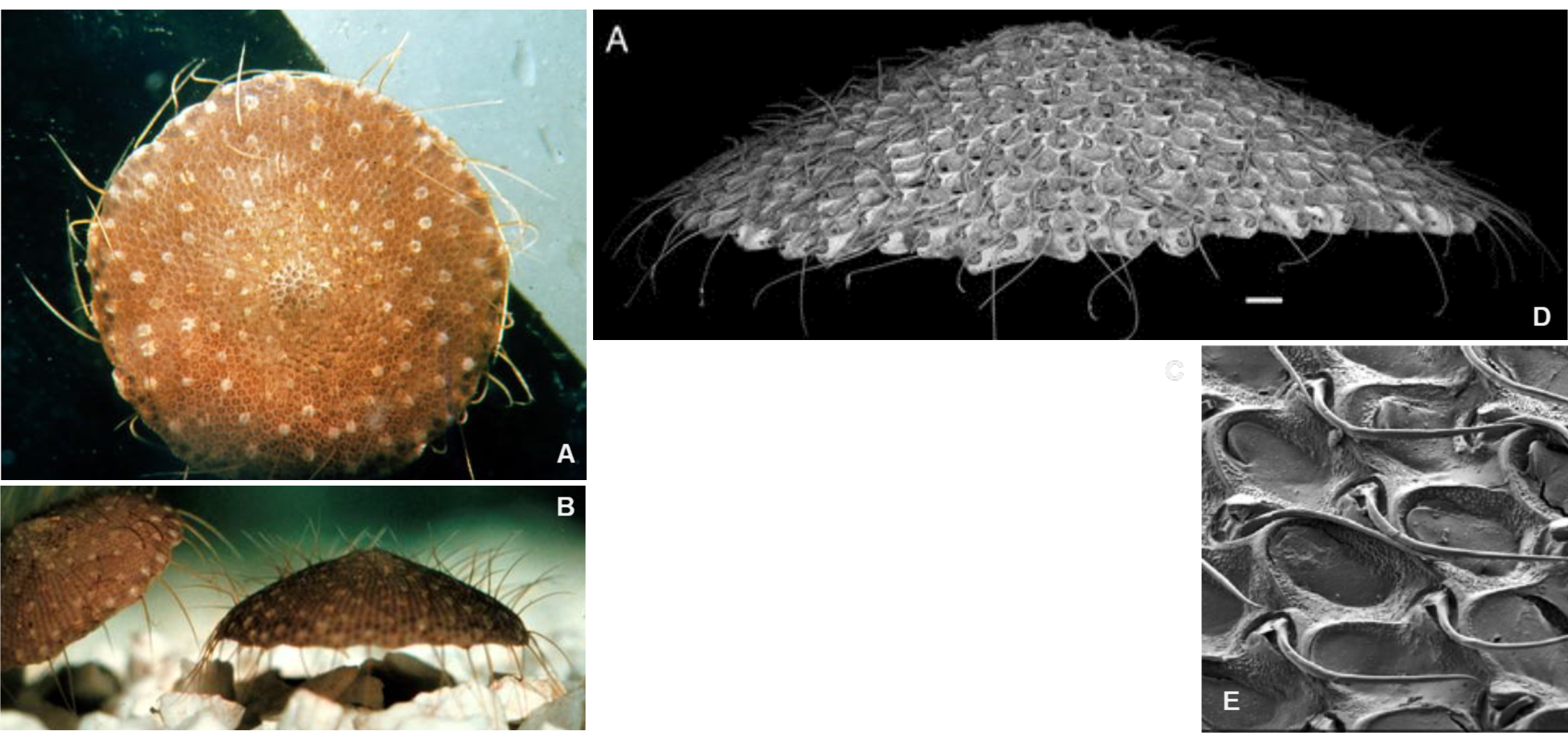
Live free-living lunulite bryozoans “walking”; note the long, flexible bristle-like setae characterizing both species. **A**-**C,** *Selenaria maculata* (Busk, 1852a), Great Barrier Reef (**A**-**B** colonies are just over 1 cm in diameter; courtesy P.L. Cook); **C**, note the low proportion of vibracula, with extremely long, serrated setae (scale bar 500 μm). **D**, *Cupuladria biporosa* (Canu & Bassler, 1923), Panama with comparatively long setae (the colony is about 12 mm in diameter). **E**, *Cupuladria guineensis* (Busk, 1854), with very short setae (courtesy P. Bock, Melbourne); note the 1:1 proportion between autozooids and vibracula with smooth setae (scale bar 200 μm).

**Fig. 2.**
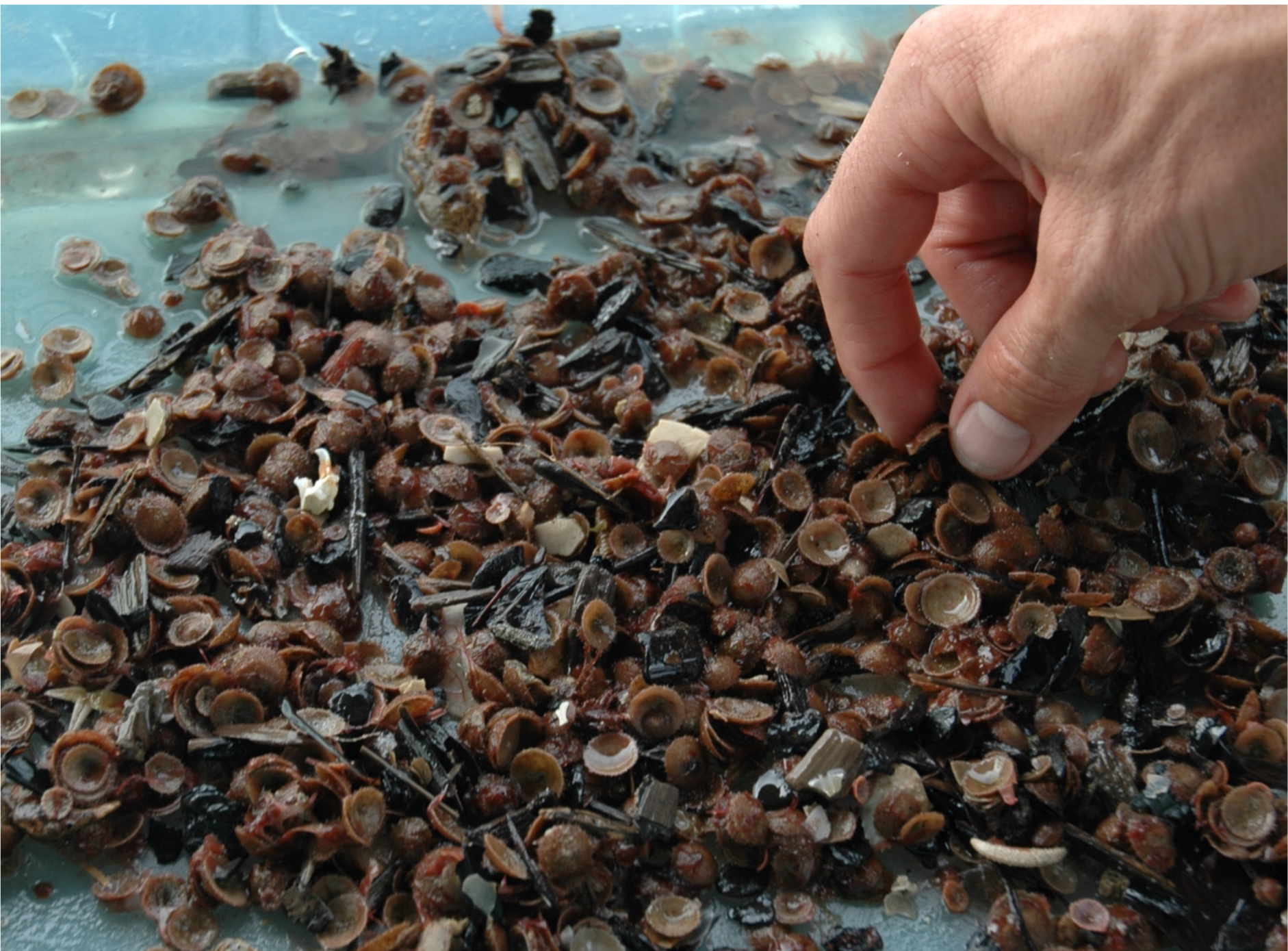
Hundreds of living colonies of *Cupuladria biporosa* collected in a van Veen grab sample in Golfo de los Mosquito, Caribbean Panama (35 m; 8.8750, −80.9992).

The free-living—or lunulitiform—bryozoans are characterized by small to fairly small (usually less than 2 cm in diameter) disc-shaped, flat to domed colonies, typically with all zooids confined to the frontal side. Their most conspicuous element is the presence of regularly distributed vibracula with individually movable long setae (Fig. 1) that extend out from the colony raising the colony slightly above the sea-floor. Representatives of two families Cupuladriidae and Selenariidae, both illustrated in Fig. 1, have been observed to move across the sea-floor, as well as digging themselves in or out of the sea-floor sediment through coordinated movement of their setae (Cook & Chimonides 1978; Håkansson & Winston 1985; O’Dea 2009) – a most unusual capability amongst bryozoans and, indeed, colonial organisms.

Bacause of the morphological constraints imposed by their mode of life, the free-living, lunulitiform bryozoans are generally easily distinguishable and, as such, a few genera were formally recognized early in the study of the bryozoans. Thus, two sets of generic names *Lunulites* Lamarck, 1816 vs. *Lunularia* Busk, 1884 and *Cupuladria* Canu & Bassler, 1919 vs. *Cupularia* de Blainville, 1830 [now abandoned], have been confused and sometimes interchanged ever since. As a result of their superficial morphological similarities, higher level classification of the groups has been ‘persistently dynamic’, with a variable number of Family-level taxa proposed, combining the lunulitiform genera in virtually all combinations possible.

Researchers have attempted to clarify the resulting systematic quagmire. In particular, a series of papers by P. L. Cook in collaboration with P. Chimonides (1978-94) and P. Bock (1998-99), among others, led to a growing realization that convergent and iterative evolution (see Box I) were likely widespread among the free-living bryozoans, a notion we support and are exploring further in a parallel investigation.

**Box I.**
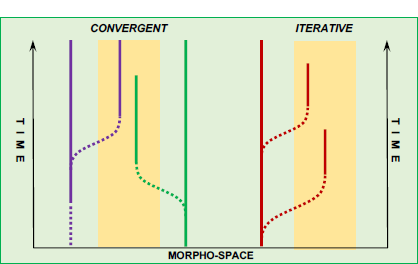
Convergent and iterative evolution. The yellow zone symbolizes the free-living morpho-space, and the line colors symbolizes separate clades.

Currently, all formally described lunulitiform taxa considered here are referred to either the Family Cupuladriidae or the Superfamily Lunulitoidea (most recently by Bock 2022), albeit commonly with some reservation. While the Cupuladriidae as a monophyletic family is well supported, we do not support the unification of all other major lunulite groups into one superfamily (see below).

## BIOGEOGRAPHY OF THE FREE-LIVING BRYOZOANS

The general pattern of free-living bryozoan biogeography is rapid occupation (and domination) of soft shelf sediments in warm waters globally following origination (Fig. 3). We illustrate the main trends in the biogeography of the free-living bryozoans by focusing on the dominant four clusters of lunulitiform free-living bryozoans, which we believe are all phylogenetically distinct, although no formal phylogenetic analyses have been made. The following summaries are based on published records and personal observations.

**Fig. 3.**
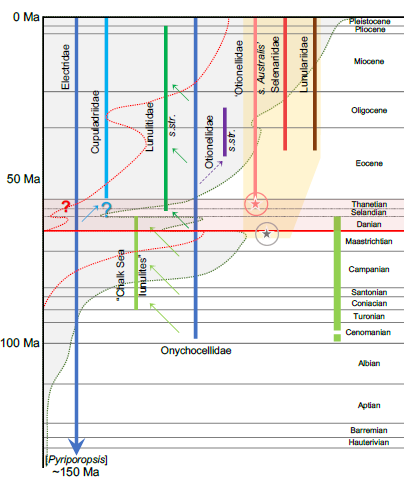
Stratigraphic distribution of the main lunulite clades referred to in the text. Two non-free-living families (dark blue lines) included as the likely ancestors to most or all of the free-living clades (indicated by thin arrows). Electridae, the longest ranging cheilostome family, include the probable ancestor to the Cupuladriidae (as well as the oldest known cheilostome genus, *Pyriporopsis* Pohowsky, 1973), while Onychocellidae is considered the likely ancestor of several ‘lunulite’ clades. Finely stippled curves indicate relative temporal diversity estimates of cheilostome [green stipple] and free-living bryozoans [red stipple], respectively (not to scale). The Paleocene “dead zone” shaded red; the “Austral Realm” shaded yellow; the temporal extent of the Northern European Chalk Sea indicated with broad green line; the two asterisks show the stratigraphic position of the two oldest known Austral lunulite records (discussed in the text).

### ’Chalk Sea lunulites‘ (Fig. 4 A, E)

This cluster of Lunulites originated in the late Cretaceous Northeast Atlantic shelf seas, most particularly the Northern European Chalk Sea, where the diversity of free-living bryozoans, from a fledgling beginning in the Late Turonian (Koromyslova & Pervushov, 2022), formed a hot-spot reaching over 100 species in the Maastrichtian. The group appears to have been geographically and ecologically constrained by the unique depositional environments of the Chalk Sea province with an assumed total temporal range from the Late Cenomanian to Danian (Håkansson et al. 1974; Surlyk 1996). The group is unequivocally <u>polyphyletic</u> (see e.g., Håkansson & Voigt 1995), with Calcitic skeletons and non-porous basal walls dominating. The group is very rare in the ultimate phase (Danian) of the Chalk Sea as well as later Paleocene deposits in the region and with unknown relations to isolated Late Cretaceous species in central Gondwana fragments (cf. Taylor 2019) and North America (undescribed). *THERE IS NO INDICATION THAT ANY MEMBER WITH RELATION TO THIS CLUSTER HAVE MIGRATED INTO THE AUSTRAL REALM*.

**Fig. 4.**
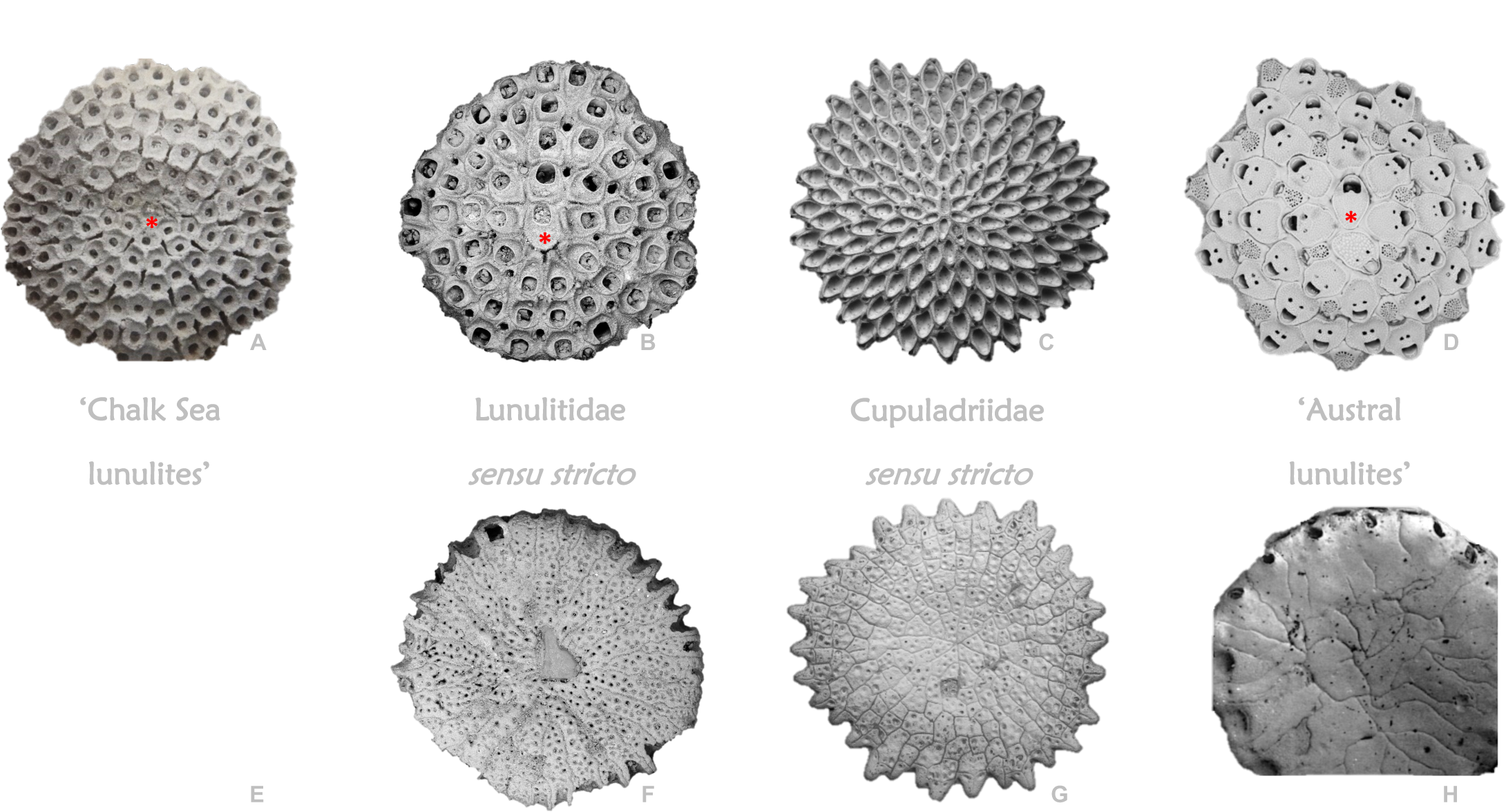
The four main clusters of lunulite bryozoans, showing both frontal (top) and basal aspects. Ancestrula and ancestrular complex indicated in red (asterisk and shaded, respectively); substrate only visible in **G** (shaded blue). **A**, *Lunulites goldfussi* von Hagenow, 1839 (Maastrichtian, Denmark). **B**, **F**, *Lunulites androsaces* Michelotti, 1838 (Late Pliocene, Altevilla, Italy). **C**, **G**, *Cupuladria biporosa* Canu & Bassler, 1923 (Recent, Florida; ancestrular triplet shaded red). **D**, *Selenaria punctata* Tennison Woods, 1880 (Recent, New South Wales). **E**, *Lunulites beisseli* Marsson, 1887 (Maastrichtian, Denmark). **H**, *Otionellina ampla* Bock & Cook, 1999 (Miocene, Victoria).

### Lunulitidae (Fig. 4 B, F)

The region of origin of this cluster as well as dominance in both abundance and diversity is the North Atlantic shelf seas, with an assumed temporal range from Selandian to Pliocene (e.g., Cipolla 1921, Lagaiij 1952; own observations). Within the Lunulitidae, two hotspots developed; a mid-Paleogene hotspot in the Northwest Atlantic region and a minor hotspot in the Neogene of the European seas. The group is probably polyphyletic, typically with a bimineralic or, less frequently, an entirely aragonitic skeleton (Taylor et al. 2009) with basal-wall pores. There appears to have been little migration outside the North Atlantic realm and, *THERE IS NO INDICATION THAT ANY MEMBER WITH RELATION TO THIS CLUSTER HAVE MIGRATED INTO THE AUSTRAL REALM*.

### Cupuladriidae (Fig. 4 C, G)

This family probably originated in the tropical eastern Atlantic (Gorodisky & Balavoine 1961). It has a known range from Thanetian or Ypresian to Recent. Apparently monophyletic (Dick et al. 2003), with dominantly aragonitic skeleton (e.g., Taylor et al. 2009), and commonly with basal-wall pores. Highly localized and rare through the Paleogene, but underwent a large-scale expansion into near circum-tropical seas at the onset of the Miocene (e.g., Laagaij 1963), forming a hotspot around the central Atlantic region. They are present, and often very abundant, in the mid-Atlantic and Caribbean, tropical west Pacific and indo-pacific archipelago from the Miocene to today, although their diversity and abundance in the latter two realms remains poorly documented. *A SINGLE CUPULADRIID SPECIES MIGRATE INTO THE AUSTRAL REALM IN THE MIOCENE* (Cook et al. 2018b).

### ‘Austral lunulites’ (Fig. 4 D, H)

With the limited evidence available, the region of origin of this cluster is most likely the Northwestern margin of the Australian continent, with a total range from Maastrichtian or Thanetian to Recent (Bock & Cook 1998, 1999; own observations). The group is polyphyletic, comprising at least four families, with dominantly aragonitic skeletons (Taylor et al. 2009). Geographically isolated through most of its history (see below), they spread around all of Australia and New Zealand by the Oligocene (Bock & Cook, 1999) and formed a prominent, ongoing hotspot with temporary migration into southern South America in the Late Oligocene, possibly *via* Antarctica. *NO AUSTRAL ‘LUNULITES’ KNOWN TO HAVE CROSSED THE WALLACE LINE INTO EURASIA*.

While the genus *Lunularia* has now been formally restricted to a small group of exclusively Austral species gathered in the monotypic family Lunulariidae (Levinsen 1909), the name *Lunulites* remains in broad use. Most particularly, it has been assigned extensively to taxa in both clusters of lunulites distinguished above, separated by not only the end-Cretaceous mass extinction, but also the subsequent Paleocene ‘dead zone’ (Fig. 3), which obscures any possible relationship between the two cohorts of *“Lunulites”.* Noteworthy, however, the scarcity of free-living bryozoans otherwise characterizing the Paleocene does not apply to the Austral realm, which was the scene of a separate, spectacular evolutionary radiation, seemingly without contribution from any lunulite clade from outside the Austral Realm (see below). As evident from the groupings above, we here advance the proposition that the two *“Lunulites*” clusters are indeed phylogenetically distinct, with “Chalk Sea lunulites” essentially becoming extinct at the Mesozoic-Cenozoic boundary, bar a few survivors into the Paleocene ‘dead zone’ (explored further by Håkansson, O’Dea & Rosso 2019 and ongoing). Together with a mixed group of unique free-living bryozoans <u>without</u> setae, *“Lunulites*” formed the – so far – unrivaled diversity hotspot of free-living bryozoans in the history of this mode of life during the later phases of the Late Cretaceous Chalk Sea in Northern Europe (e.g., Håkansson & Voigt 1995).

## THE ‘AUSTRAL LUNULITES’

The biogeographic realm encompassing Australia and New Zealand is traditionally referred to as Australasia (e.g., Gordon et al. 2019); however, in the framework discussed here this term is manifestly illogical and misleading, since a crucial element in the concepts presented here is the strict biogeographic separation between Australia and Asia – well established for the terrestrial biota but equally relevant for marine faunas, bryozoans included.

Consequently, we here refer to this province as the ‘Austral Realm’. This realm occupies a prominent position in the study of Cenozoic and modern bryozoans – including the lunulites – with highly diverse faunas being extensively studied since the early 19^th^ century. Initially based, albeit rather chaotic, on the collections from the French Baudin expedition (1801-03); but following the British “Rattlesnake” expedition (1846-50) and subsequent locally instigated collecting, a more systematic description of the bryozoan faunas commenced (e.g. Busk 1852a,b, 1854; Hincks 1881a,b, 1882; MacGillivray 1881, 1886; see Cook et al. 2018a for an overview). Among the earliest work on fossil bryozoans are Tenison Woods (1865) as well as a monograph by MacGillivray (1885), with early works on fossil lunulite bryozoans by Tenison Woods (1880) and Maplestone (1904), all referred to the Family Selenariidae (but see below).

As is evident from the groupings outlined above, we maintain that the Austral free-living bryozoans remained largely isolated from the rest of the World throughout most of their history, a proposition hinted at by previous authors (e.g., Cook & Chimonides 1985b; Bock & Cook 1998). While this proposition was previously based primarily on ancient and modern distributions of the different groups, it is also consistent with the post-Gondwanan history of isolation of Australia (Fig. 5). By the time the first free-living cheilostome bryozoans evolved in Eurasia (latest Turonian), only Antarctica [with New Zealand] was still attached to Australia, forming a southern continent surrounded by substantial oceans on most sides as well as a cold-water barrier across Antarctica and into South America (see e.g., Müller et al. 2021). Late Paleogene separation between these three terranes sent Australia and New Zealand on somewhat different northward trajectories which would, eventually, lead towards the post-Miocene closure of the Tethyan Ocean north of Australia, with only a few, narrow deep-water straits across the Indonesian archipelago remaining (e.g., Barber et al. 2000). In short, not before the Middle Eocene did these terranes reach a position facilitating at least some warm, shallow water connections to the rest of the World (*in casu* SE Asia, as evidenced by the first occurrence of warm-water benthic foraminifera in Australia; Haig et al. 1997). This prolonged isolation of the Austral shallow marine realm, commencing well before the appearance of the first lunulite bryozoan in Europe, therefore supports the notion that the ‘Austral lunulites’ evolved their free-living mode of life independently from any outside group. In all probability it therefore provides an independent parallel to the simultaneous evolution of traits particular to the free-living mode of life in the Lunulitidae s. str. and the Cupuladriidae (Håkansson et al. 2019).

**Fig. 5.**
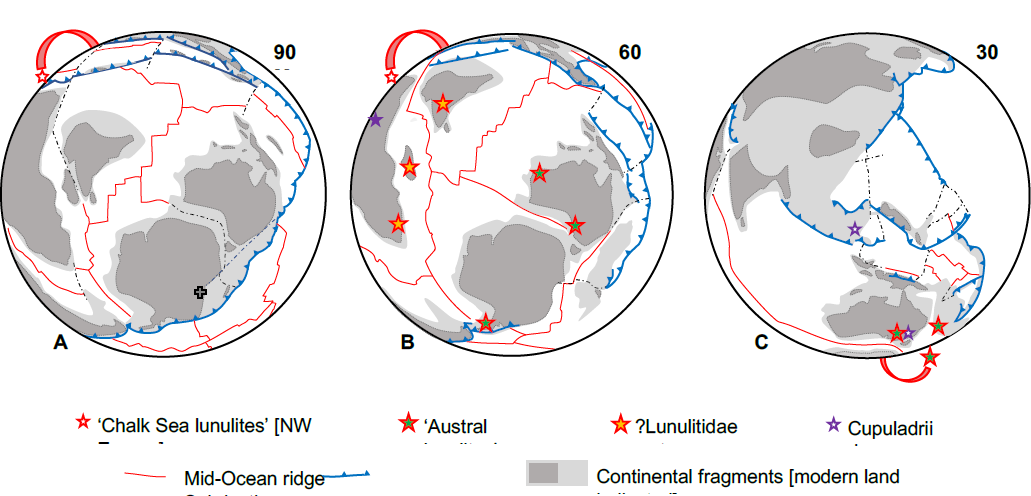
Consecutive plate configurations (simplified from Müller et al. 2021) as background for the distribution of lunulite bryozoans from their first appearance in the Late Turonian, in the European Chalk Sea, through to the early Neogene. **A**, The early part of the Chalk Sea; few lunulites and only in Northern Europe and Asia. The rough position of the 90 Ma South Pole indicated (with a cross), with only limited subsequent change in position. **B**, Encompassing the rapid change across the Mesozoic-Cenozoic transition (including data for the period Maastrichtian - Early Eocene). **C**, Encompassing the comparative short period with invasion of Austral lunulites into southern South America (Late Oligocene - Middle Miocene). Note the continuous isolation of the Australia-Antarctica-New Zealand-southern South America remnants of Gondwana locked in high latitudes traditionally considered unsuited for lunulite bryozoans.

The origin of the distinct Austral brand of free-living bryozoans remains somewhat ambiguous due to a very limited early fossil record. The Late Aptian record of *Lunulites abnormalis* Etheridge, 1901 was refuted by Håkansson et al. (in press) in the absence of the original and only known specimen combined with woefully inadequate description and illustrations. On this background, the potentially earliest known appearance is a less dubious record from the latest Maastrichtian Miria Marl formation in Western Australia reported by Cook & Chimonides (1986), exposed in the now onshore part of the Southern Carnarvon Basin. The earliest substantiated occurrence is therefore a fauna comprising at least 10 species level taxa from the Thanetian Boongerooda Greensand (EH collections; see further below). However, Already by Late Eocene all modern families of lunulite bryozoans in the Austral realm were present in the significant expansion of Austral free-livings, with dominance and diversity peaks shifting to southeastern Australia and, in the Oligocene, to include New Zealand, forming an independent Austral lunulite hotspot reaching its diversity maximum during the Miocene (Bock & Cook 1999). Around the same period, the Austral free-living fauna experienced a brief interval of extra-continental expansion into southern South America, possibly facilitated by a more favorable Late Oligocene – Middle Miocene temperature regime (see below).

Even though it appears unlikely that any Austral free-living taxon has a free-living ancestral connection outside the Austral region [the apparently Miocene immigrant *Cupuladria guineensis* Busk, 1854 being the exception], we suggest it probable that the group is polyphyletic, strengthening the conclusion that becoming a free-living bryozoan may be relatively easily achieved. We currently consider the following family level clades to have originated in the Austral realm: Lunulariidae, Selenariidae, and ‘Otionellidae’ (amended, see below), all well known throughout the Late Eocene to Recent (see Bock & Cook 1998, 1999 & references therein). Ongoing work on the undescribed faunas from the Thanetian and Maastrichtian(?) in Western Australia will add at least two new families from the Thanetian (presented briefly below) and – potentially – an additional new family from the Maastrichtian of the same region.

### Lunulariidae Levinsen, 1909

The monogenetic family Lunulariidae Levinsen, 1909 (Figs 6 & 8 F) is known from the Late Eocene to the present in Australia and New Zealand (from the Miocene onwards; Bock & Cook 1998). Female zooids (barely recognizable from the exterior in some species) with expanded volume to accommodate the ovisac, are brooding a mega, presumably nutrient packed larva (Cook & Chimonides, 1986) allowing prolonged metamorphosis resulting in a large ancestrular complex comprising up to 12 autozooids (ordinary feeding zooids) in *Lunularia capulus* before feeding commences (EH unpublished data, *in* Cook & Chimonides 1986). Despite several clear statements to the contrary (e.g., Bone & James, 1993), the ancestrular complex is consistently without a known substrate (EH unpublished paper, read at the Vienna 1981 IBA Conference; Cook & Chimonides, 1986; Cook et al. 2018b).

**Fig. 6.**
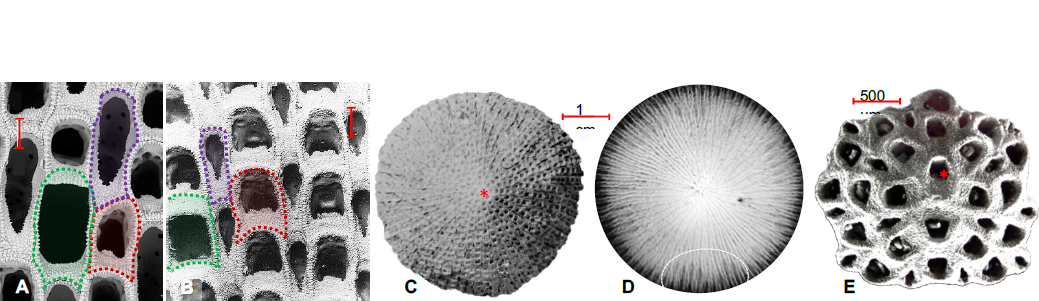
Selected species of Lunulariidae. **A**, *Lunularia repanda* Maplestone, 1904; Investigator Strait, South Australia, Recent. Detail showing zooidal morphology, note the significant size increase in female zooids. **B**, *L. capulus* (Busk, 1852a); Recent; detail showing zooidal morphology; note that female zooids are externally recognizable only by their slightly enlarged, square opesia. **C**, *L. capulus* (Busk, 1852a); Recent; frontal side of small, mature colony with perfect radiating symmetry (ancestrula marked with red asterisk), and **D**, x-ray image of the same colony revealing the distribution of the internally enlarged female zooids (diffuse, light grey shades) in incomplete circles near the margin of the colony; one cluster indicated by thin white stipples. Note the complete lack of a substrate. **E**, *L. capulus* (Busk, 1852a); (SAM7423, Recent; Juvenile colony. Zooidal color-codes (Figs 6-7 & 11): autozooids red, vibracula purple, female zooids (ovicells) green, male zooids blue). Scale bars: **A**-**B** 200 μm. (**A-B** courtesy Phil Bock, Melbourne, **E** courtesy Mary Spencer-Jones, NHM London).

**Fig. 8.**
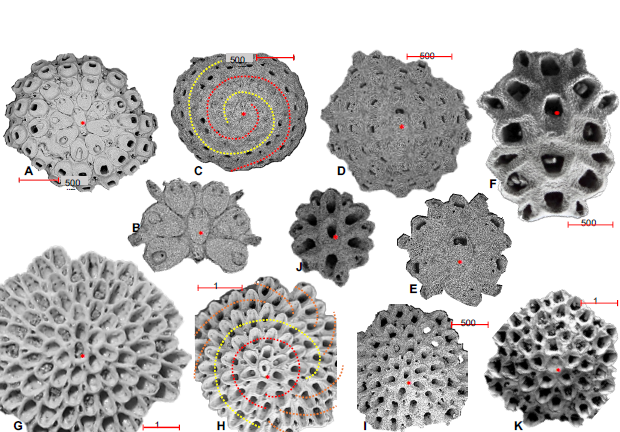
Colony architecture and growth pattern in selected free-living Austral species (ancestrula indicated with red asterisk). **A**-**B**, *Otionellina cupula* (Tennison Woods, 1880); Victoria, Miocene. Frontal side of complete colony, with the ancestrular complex magnified (**B**); note closure of the central zooids. **C**, *Helixotionella spiralis* (Chapman, 1913); Victoria, Miocene). Frontal side of complete colony, with the two spiral budding rows indicated (red and yellow stipple). **D**-**E**, *Selenaria punctata* Tennison Woods, 1880; New South Wales, Recent. Frontal side of complete colony, with the ancestrular complex magnified (**E**). **F**, Ancestrular complex of *Lunularia capulus* (Busk, 1852a); SAM7423, Recent; cut-out from Fig. 6E. **G**, *Discoradius*(?) *rutella* (Tenison Woods, 1880); Victoria, Miocene. Complete colony with radiating budding; note the 4 vibracula in association with the ancestrula. **H**, *Kausiaria magna* Bock & Cook, 1998; Victoria, Late Eocene; frontal side of complete colony with the complex spiral growth pattern indicated, with the ancestrular complex magnified (**E**). **I-J**, *K. jamesi* Bock & Cook, 1998; Victoria, Late Eocene. Complete colony showing the more traditional radial budding pattern. **K**, complete colony of *Petasosella lata* (Tennison Woods, 1880); Victoria, Miocene. (Images courtesy Phil Bock, Melbourne, except **J**, courtesy Mary Spencer-Jones, NHM, London, and **H**).

*Lunularia* Busk, 1884 – In many ways this genus is close to the genus *Lunulites*, with many morphological features resembling members of both the ‘Chalk Sea lunulites’ and the Lunulitidae. On the other hand, the presence of an interior ovisac seemingly parallels the reproductive life history strategy of the Cupuladriidae (cf. Ostrovsky et al. 2009; Ostrovsky 2013; and references therein). Members of this genus form some of the largest known free-living colonies, reaching diameters over 7 cm, as estimated from the size of fragmented colonies (EH data, from the Geology collections at University of Southern Australia, Adelaide). The ancestrula is comparable in size to the periancestrular autozooids, and there are <u>no</u> vibracula in direct contact with the ancestrula.

### Selenariidae Busk, 1854

The monogenetic family Selenariidae Busk, 1854 (Figs 7 A-H & 8 D, E) is known from the Late Eocene to the present, with a significant number of species from Australia, a single, recent species reported from New Zealand (Bock & Cook 1999), and at least two species from the southern part of Argentina and Chile (Late Oligocene and Middle Miocene; López-Gappa et al. 2017; EH collections). The family is characterized by advanced sexual polymorphism with distinct male and female zooids in many taxa (Cook & Chimonides 1985a, 1985b, 1987; Bock & Cook 1999 & references therein).

**Fig. 7.**
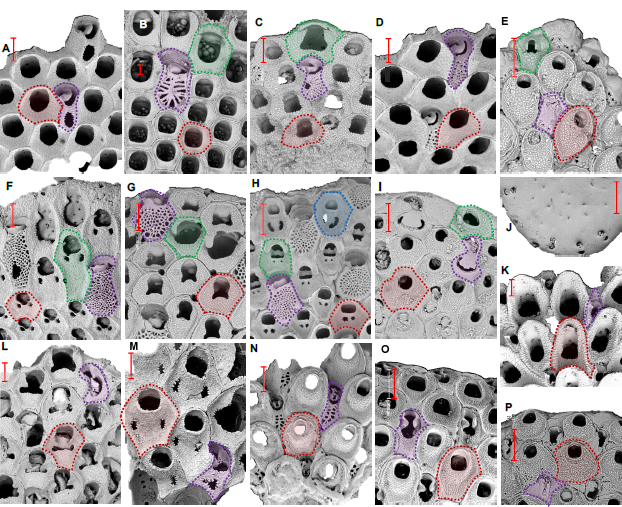
Selected Recent and fossil species of Selenariidae (**A**-**H**) and ‘Otionellidae *sensu Australis’* (**I**-**P**) illustrating the range of zooidal polymorphism characterizing these families: two to four types of zooids in the Selenariidae, and two to three types in the ‘Otionellidae’. **A**, *Selenaria concinna* Tennison Woods, 1880; Victoria, Recent. **B**, *S. hexagonalis* Maplestone, 1904, Investigator Strait, state, Recent. **C**, *S. verconis* Parker & Cook, 1994, Victoria, Miocene. **D**, *S. kompseia* Cook & Chimonides, 1987; New South Wales, Recent. **E**, *S. initia* (Waters, 1883); Victoria, Miocene. **F**, *S. bimorphocella,* Mapelstone, 1904; South Australia, Recent. **G**, *S. minor* Maplestone, 1911; Victoria, Recent. **H**, *S. punctata* Tennison Woods, 1880; Recent. **I**-**J**, *Helixotionella scutata* Cook & Chimonides, 1984b, Western Australia, Recent. **K**, *Kausaria magna* Bock & Cook, 1998; Victoria, Late Eocene. **L**, *Petasosella alata* Tennison Woods, 1880; Victoria, Miocene. **M**, *P. magnipunctata* (Maplestone, 1904); Victoria, Miocene. **N**, *Otionellina squamosa* (Tennison Woods, 1880); New Zealand, Pleistocene. **O**, *O. australis,* Cook & Chimonides, 1985; Victoria, Recent. **P**, *H. spiralis,* (Chapman, 1913), Victoria, Miocene. (Scale bars 200 μm; zooidal color-codes as in Fig. 6. Images courtesy Phil Bock, Melbourne).

*Selenaria* Busk, 1854 – The genus is characterized by the zonal distribution of sexual polymorphs and large, scattered, often very complex vibracula (proportionally significantly fewer than autozooids), with long setae observed to facilitate locomotion in several species (Cook & Chimonides 1978). The frontal walls of both autozooids and vibracula are highly variable. The ancestrula is comparable in size to the periancestrular autozooids, and there are typically <u>no</u> vibracula in direct contact with the ancestrula (Fig. 8 E).

### ‘Otionellidae sensu Australis’

In the current understanding, the family Otionellidae Bock & Cook, 1998 (Figs 7 I-P & 8 A-C, H-K) comprises the following 5 genera: the nominate genus *Otionella* Canu & Bassler, 1917 in addition to *Otionellina* Bock & Cook, 1998, *Helixotionella* Cook & Chimonides, 1984b, *Petasosella* Bock & Cook, 1998, and *Kausiaria* Bock & Cook, 1998. However, while the nominal genus of the family Otionellidae only accommodates a few closely related species from the Middle Eocene to Early Oligocene in North America, all other genera have a distinct Austral distribution. In our opinion, and already indicated by Bock & Cook (1998, e.g., their p. 197 “….a very ancient southerly group of lunulitiform bryozoans.”), biogeography as well as pronounced morphological differences suggest a separation between a restricted, North American family Otionellidae *s. str*., and a strictly Austral – as of yet – unnamed family, here provisionally referred to as ‘Otionellidae *sensu Australis’* (in prep.). It comprises at least four genera, three of which were temporarily present also around southern South America (Canu 1904; Bock & Cook 1998; Pérez et al. 2015; EH collections). Based on the distribution of the periancestrular autozooids and vibracula, these genera may be further subdivided into two groups – one with<u>out</u> vibracula in direct association with the ancestrula (*Petasosella* and *Kausaria*), and the other with a distolateral and a proximal vibraculum in direct association with the ancestrula (*Otionellina* and *Helixotionella*). Based on morphological similarities in the skeletal structure of the vibracula, as well as the presence of sexual polymorphism (Fig. 7), it appears possible that the second entity of this geographically restricted and currently unnamed family may share a common ancestry with the Selenariidae.

*Petasosella* Bock & Cook, 1998 (Figs 7 L-M & 8 K) – This genus is characterized by a radiating budding pattern with an ancestrula surrounded only by autozooids. The vibracula are large and scattered, proportionally fewer than autozooids. No skeletal sexual polymorphism noted and, therefore, with potential brooding in interior ovisacs. The genus is known from the Late Eocene to Recent of southeastern Australia, and – plausibly – Late Oligocene to Early Miocene in southernmost Argentina (Canu 1904; Bock & Cook 1998; EH collections).

*Kausiaria* Bock & Cook, 1998 (Figs 7 K & 8 H-J) – This genus is characterized by comparatively large colonies, an ancestrula surrounded exclusively by autozooids, and a variable budding pattern – either radial *or* composite sinistral spirals (Figs 8 D and I, respectively). The vibracula are small and proportionally equaling autozooids. Colonies consistently without a substrate. The genus is known from the Late Eocene to Miocene of southeastern Australia, and from the Late Oligocene of southernmost Argentina (EH collections).

*Otionellina* Bock & Cook, 1998 (Figs 7 N-O, 8 A-B) – This genus is characterized by its compact, almost lenticular colonies, with abundant, fairly small vibracula (proportionally equaling autozooids), which are commonly developed also along the margin of the basal side of the colony. The ancestrula characteristically has one distal and one proximal vibraculum (Fig. 8 B). The genus – or close relatives - is known from the Thanetian of Northwestern Western Australia (see below), from the Middle Eocene – Recent in southeastern Australia (Bock & Cook 1998, 1999; Schmidt 2007), from the Oligocene – Recent in New Zealand (Bock & Cook, 1999), and from the Late Oligocene to Middle Miocene in southern South America (Canu 1904; Pérez et al. 2015; EH collections), as well as – possibly – from the Early Eocene in Antarctica (Hara et al. 2018; their fig. 7, <u>only</u>).

*Helixotionella* Cook & Chimonides, 1984b (Figs 7 I-J, 9 C) – This genus is characterized by its unusually small, typically lentil-shaped colonies, abundant small vibracula (proportionally equaling autozooids) and a spiral budding pattern, with two distinct, dextral budding series. The budding series are initiated from the two vibracula associated with the ancestrula. In late astogeny the budding series may bifurcate, and all or most budding series are associated with a (terminal) vibraculum on the basal side of the colony, strongly suggesting terminate growth of the colonies. The genus is known from the Late Eocene to Recent in Southeastern Australia, with a potential relative in the Thanetian of Northwestern Australia (see below)

### Incertae sedis

Together with its nominal family Lunulitidae Lagaaij, 1952 the genus *Lunulites* Lamarck, 1816 is currently very loosely constrained and, in the tradition outlined above, it has been variously applied also to Austral free-living taxa. Most have subsequently been referred to the endemic Austral genera listed above, but one fossil taxon – quite widespread in the Australian Paleogene and Neogene successions – remains largely unaccounted for in terms of its taxonomic and – indeed − phylogenetic affiliation. Originally described as *Lunulites rutella* Tenison Woods, 1880 (our Fig. 8 G), this species has fairly consistently been referred to as *Lunulites*, albeit commonly with some reservation (see Bock & Cook 1999 & references therein), and recently it has been transferred to the new genus *Discoradius* Di Martino, Greene & Taylor, 2017, again with some reservation (Di Martino et al. 2017). In our opinion, the zooidal morphology suggests a closer relationship to the austral genus *Kausiaria*. However, the quite unique early astogeny, with three [occasionally four] small periancestrular vibracula (a disto-lateral pair of vibracula directed proximally plus one [or two] proximal vibracula), may suggest yet another independent free-living Austral clade.

## ORIGIN OF THE AUSTRAL LUNULITE HOTSPOT

Considering the known distribution of fossil free-living bryozoans within the Austral Realm (comprehensive summaries in Bock & Cook, 1998, 1999, supplemented by EH collections from South America and Northwestern Australia), the origin of this evolutionary isolated lunulite cluster was likely the Northwestern margin of the Australian continent, in the proximal part of the broad, gently sloping shelf bordering the Indian Ocean in a configuration similar to the present. Reflecting its very proximal position on this platform, the stratigraphic succession containing these faunas is significantly condensed with sediment accumulation restricted to transgression maxima (Fig. 9). Thus, the two main intervals from where free-living bryozoans have been reported – the Late Maastrichtian Miria Marl Formation (Cook & Chimonides 1986; two loose specimens) and the immediately overlying Thanetian Boongerooda Greensand Member (EH collections), initiating the Thanetian-Ypresian Cardabia Formation, are both thin [up to a couple of meters], but separated by a significant lacuna comprising the Mesozoic-Cenozoic boundary interval (Fig. 9).

**Fig. 9.**
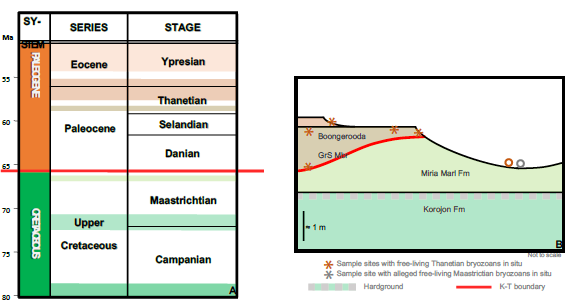
Stratigraphy of the Late Cretaceous - Early Paleogene succession in the on-shore part of the Southern Carnarvon basin on the Northwestern margin of the Australian continent. **A**, Chronostratigraphic chart covering the Upper Cretaceous and Paleogene strata in the Giralia Anticline. Note the frequent and extensive breaks in accumulation reflecting the position of this succession in the proximal part of a passive continental margin. **B**, Highly stylized cross section illustrating the field relations of the strata investigated. The levels yielding free-living bryozoans in situ marked by asterisks, while the location of loose specimens is indicated with a circle – alleged Late Maastrichtian grey, Thanetian brown. Note that the only two specimens referred to the Maastrichtian are collected loose. (The level of the end-Cretaceous mass-extinction event marked by red line. GrS Mbr= Greensand Member; Fm = Formation).

The two specimens of lunulite morphology referred to as originating from the Maastrichtian Miria Marl formation share a dubious provenance. According to their labels they were collected from loose material but assigned as being from the Miria Marl section (WAM, 80.648, 80.649, Miria Marl Fm, Cardabia Station; Cook & Chimonides 1986; our Fig. 10). However, there is no adhering sediment to support this assignment and, as seen from Fig. 9, it is equally plausible that they originated from the younger Boongerooda Greensand Member and were simply mislabeled. Both specimens are unequivocally free-living, but are poorly preserved, allowing only a limited amount of certainty with respect to their taxonomic position. One, WAM 80.649 (Fig. 10 A-C), may be an early relative of *Otionellina* (cf. Figs 7 &, 8), but WAM 80.648 (our Fig. 10 D-G) does not resemble any of the free-living taxa in the overlying Boongerooda Greensand, suggesting it *could* be Maastrichtian in age. Moreover, the moderately rich free-living fauna from the Thanetian Boongerooda Greensand is reasonably well preserved, and has clear ties with the later, better-known Austral free-living faunas – most obviously, but not exclusively, with the family ‘Otionellidae *sensu Australis’.* For the present, we leave the taxonomic as well as the stratigraphic status of the two specimens referred to the Miria Marl open, pending further investigation.

**Fig. 10.**
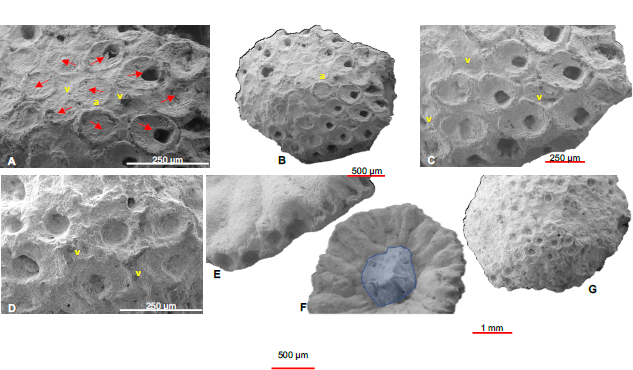
Possibly the oldest lunulite bryozoans from Australia, collected loose and for untold reasons referred to the Late Maastrichtian Miria Marl (see text). **A**-**C**, Early, potential representative of ‘Otionellidae’ *sensu Australis;* WAM 80.649. **A**. The periancestrular region with arrows indicating the interpreted orientation of the ancestrula (a) and the early autozooids; two potential vibracular marked (v). **B**. Overview, centered around the best-preserved part of the frontal surface; ancestrula indicated. **C**. Colony margin with several well-preserved autozooids and scattered, variously preserved vibracula (v). **D**-**G**, Indeterminate lunulite colony; WAM 80.648. **D**. The ‘best’ preserved part of the frontal side providing enough detail of the autozooidal morphology to exclude the likeliness of conspecificity with WAM 80.649; potential vibracula indicated (v). **E**. Oblique basal view of the well-preserved colony margin. **F**. Basal surface of the colony displaying distinct budding rows with secondarily thickened calcitic walls and widely scattered minute pits, which may represent basal pores; note the sand-size substrate (shaded blue). **G**. Overview, showing the seriously corroded frontal side of the colony

### New family X (in prep.; Fig. 11 A-G)

This family is thus far known only from the Thanetian and possibly Ypresian of the Southern Carnarvon Basin. Members referable to this family are interpreted to be free-living <u>without</u> setal support, and without a substrate, a condition which is otherwise frequent only in relation to the Late Cretaceous, Northern European Chalk Sea (Håkansson 1975). However, the Austral version of this variety of free-living is unlike anything seen anywhere else, comprising perhaps four different species, all sharing essentially identical zooid morphology, but with significant differences in colony architecture. Similar to most free-living bryozoans from the Chalk Sea <u>without</u> setal support, the Austral species are presumably going through metamorphosis without the physical support of a substrate. To what extent the considerable architectural variation encountered within this new family warrants the recognition of more than a single genus is still under consideration.

**Fig. 11.**
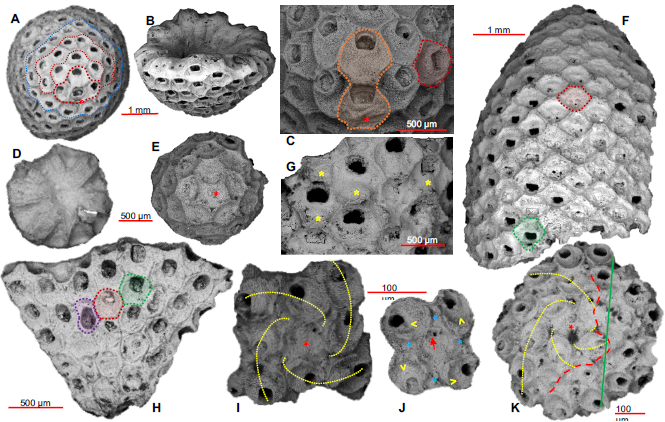
Selected free-living taxa from the Thanetian of the Southern Carnarvon Basin, Northwestern Western Australia. **A**-**G**, New Family **X**; two unnamed taxa illustrating the morphological range of the family. **A**-**C**, New species **a. A**, Mature colony; the early fan-shaped growth-stages outlined in red stipples (ancestrula with red asterisk; ancestrular complex with thick stipple; subsequent stages with thin stipple) and the first, subsequent stage with radial budding outlined in blue stipples. **B**, Oblique lateral view of the same colony showing the hollow, hemispherical shape. **C**, Detail of the ancestrular region of the same colony showing the single, ovicell complex (orange shading) developed from the ancestrula (red asterisk) late in astogeny, through partial skeletal resorption in the distal autozooid. **D**-**G**, New species **b**. **D**, Basal view of juvenile, starshaped colony demonstrating the symmetric budding pattern radiating from the ancestrula. **E**, frontal view of juvenile colony, maintaining the strict radial budding pattern from the central ancestrula (red asterisk). **F**, Lateral view of a large fragment of a mature colony demonstrating the highly unusual, very tall, to near cylindrical colony shape. **G**, Detail of the mature part of the same colony with several ovicells/female zooids (yellow asterisks mark the low arching ovicells). **H**-**K**, Two un-named taxa referred to the provisionally named ‘Otionellidae’ *sensu Australis.* **H**, cf. *Otionellina* n. sp., fragment of discoidal colony with radial budding showing zooidal details. **I**-**K**, aff. *Helixotionella* n.sp. **I**, Complete juvenile colony comprising the first two zooids in each of the four sinistral spiral budding rows (ancestrula indicated with red asterisk, budding rows in yellow stipples). **J**, Cut-away detail of **I** showing the distribution and relative orientation of the ancestrula (red arrow), the initial four autozooids (yellow <, indicating orientation), and the four primary vibracula (blue asterisks). **K**, Complete, regenerated colony, with spiral budding (yellow stipples) in the original part of the colony (including the ancestrula, red asterisk) and chaotic budding in the regenerated part (fracture line in red stipples). Additional zooidal color-code as in Fig. 6.

Two members of this new clade are illustrated here. New species *a* (Fig. 11 A-C) is characterized by small (up to c. 4 mm in diameter) domed to hemispherical colonies with a consistent budding pattern, characterized by an early stage with gradually expanding fan-shaped budding from a marginal ancestrula which, after three budding generations, changes into a radial budding pattern gradually bringing the ancestrula into a more and more central position (cf. Fig. 11 A). When mature, a <u>single</u> ovicell (brood chamber) may develop from the ancestrula, through partial skeletal resorption also involving the distal zooid (Fig. 11 C), in a process and position that is unknown anywhere else in the phylum. In contrast, new species *b* (Figs 11 D-G) has a budding pattern with a central ancestrula from the onset (Fig. 11 E) and a subsequent, perfectly radial budding symmetry (Fig. 11 D) rapidly turning the colony into a tall (>1 cm), hollow column with near parallel sides (Fig. 11 F), maintained through a slow increase in the number of budding series. The ancestrula consistently occupies its central position at the apex of the column and, unlike species *a*, this taxon develops multiple ovicells in female zooids in a more ‘normal’ position, several budding generations away from the ancestrula (Fig. 11 G).

The remaining, less abundant, free-living taxa from the Boongerooda Greensand comprises a fair number of apparently not very closely related taxa, all (potentially) possessing vibracula. The early history of free-living bryozoans in Western Australia is based entirely on taxa yet to be formally described (in prep.) although it is already evident that some of these pioneers may indeed be directly related to well-established entities in the later Austral faunas, with two such examples illustrated here (Fig. 11 H-K).

aff. *Otionellina* n. gen., n. sp. (in prep.; Fig. 11 H) has flat, disc-shaped colonies with zooids confined to one side, budding is radial, and it resembles *Otionellina* in zooid morphology and being without a substrate; however, the thick, lenticular shape and basal side vibracular characteristic of that genus, are not present in this taxon. The autozooids have a depressed, granulated cryptocyst with a large, terminal opesium; ovicells are recognized through the low, hood-like structure distal to the opesium combined with a more angular, almost quadratic outline of the opesium. The vibracula are fairly large, typically initiating budding rows; they are mostly slightly asymmetrical, with variously developed condyles and – pending information on the early astogeny – this taxon may either represent an early member of the genus *Otionellina* or a new genus related to *Otionellina*.

aff. *Helixotionella* n. gen., n. sp. (in prep.; Figs 11 I-K). This taxon is characterized by its miniscule colonies (mostly <1 mm) with abundant vibracula (proportionally equaling autozooids in number) and a spiral budding pattern with *<u>four</u>* sinistral budding series, each initiated from one of the <u>four</u> vibracula associated with the ancestrula. The genus is known only from the Thanetian of Northwestern Western Australia, and provisionally we consider this new genus a relative of, and potential ancestor to *Helixotionella*.

## THE SOUTH AMERICAN CONNECTION

Prior to the Late Oligocene, free-living, lunulite bryozoans have a poor record in South America comprising just a single Paleocene species of *Discoradius* from Northeastern Brazil (Buge & Muniz 1974; Di Martino et al. 2017, 2018). However, during the Late Oligocene to Middle Miocene, a moderately diverse cupuladriid fauna coexisted with the Austral emigrants around the southern part of the continent (Fig. 12 A), commencing at a time where the Cupuladriidae were very rare in the rest of the World. However, both groups are today absent from this part of the continent, probably because of considerable late Miocene to Pleistocene cooling.

**Fig. 12.**
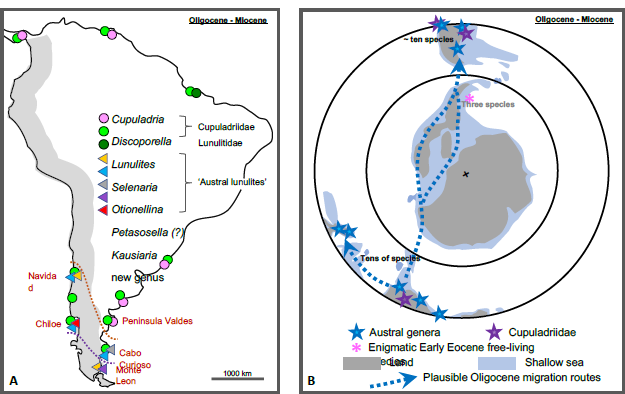
**A**, Distribution of free-living bryozoan taxa in South America during the Late Oligocene - Middle Miocene as presently known. Note the presence of cupuladriids across the continent (except the most extreme south) overlapping significantly with the Austral genera in the southern part (Canu 1904, 1908; Pérez et al. 2015; López-Gappa et al. 2017; own observations). The apparent absence of Cupuladriids between the documented presence of Oligo-Miocene cupuladriids in Darian, Panama (ref) and Navidad, Chile reported here, is likely to reflect insufficient sampling. However, the effect of cold upwelling along the west coast of South America somewhat comparable to the modern situation cannot be discarded. **B**, Paleogeographic map showing a plausible migration route connecting the Austral realm to South America via Antarctica; note that all cupuladriid taxa on either side of Antarctica arrived from the North, with <u>no</u> connections across the polar Antarctic seas. Note also that while there is a significant overlap at the genus level between the Austral realm and southern South America, the three Early Eocene lunulite taxa from Antarctica (Hara et al. 2018) do not appear closely related to either of these regions (see text). The paleogeographic outline in **B** based on Lawver & Gahagan (2003).

The presence of an Austral lunulite fauna in Upper Oligocene to Middle Miocene strata in southern Argentina and Chile may be summarized as follows (Fig. 12 A). Approximately ten taxa with Austral connections have been distinguished: three species referable to *Selenaria*, one species referable to *Otionellina*, two species closely associated with *Petasosella*, one species referable to *Kausiaria*, as well as two species that may warrant the creation of new genera, one related to *Selenaria* and one possibly related to *Otionellina* (Phillipi 1887; Canu 1904, 1908; Pérez et a al. 2015; López-Gappa et al. 2017; supplemented by EH data). For all genera listed, these species represent the only record outside their Austral heartland, thus overwhelmingly pointing to a close faunal relation between the Austral and Magallanes biogeographic provinces during this period.

The existence of a significant Late Oligocene to Miocene marine connection between the Austral realm and the South American Magallanes province have been pointed out previously, particularly based on various Mollusca that were thought to have dispersed in the circum-Antarctic current system (e.g., Beu et al. 1997). However, while the time of deep-water opening between Tasmania and Antarctica is fairly well established to be about the Eocene-Oligocene boundary, the Drake Passage between Antarctica and South America is less so, with estimates ranging from late Eocene to early Miocene (Scher & Martin 2006 Dalziel et al. 2013, Hodel et al. 2021). Nevertheless, as pointed out by Casadio et al. (2010), declining faunal similarities between the Austral and Magallanes provinces occurred *after* the establishment of the circum-Antarctic current and, therefore, an alternative, shorter route of dispersal *via* the archipelago formed in the West Antarctic Rift System would seem more likely (Fig. 12 B). And considering the assumed short-lived larval stages of modern lunulite taxa, this alternative would indeed seem more attractive.

Three lunulite bryozoans recently described from the mid Ypresian of the Antarctic Peninsula (Hara et al. 2018) offer some support to the existence of this connection, although just a single species - *Otionellina antarctica* Hara, Mörs, Hagström & Reguero, 2018 – is potentially showing any relation to the core Austral province. Nevertheless, the mere presence of a free-living Antarctic fauna clearly demonstrates that suitable habitats were present, hence providing supporting evidence for a trans-Antarctic route of migration between the Austral Realm and South America along the West Antarctica archipelago (Casadio et al. 2010; our Fig. 12 B). To further complicate comparisons, it should be emphasized that the Antarctic fauna of Early Eocene age is older than both neighboring faunas and that it shows even less resemblance to the new Late Paleocene – earliest Eocene faunas from Northwestern Australia.

## GLOBAL FREE-LIVING BIOGEOGRAPHY – AN OVERVIEW

The biogeography of the lunulite bryozoans has been a matter of interest over the years, with a series of distributional maps presented by Cook & Chimonides (1983, their Figs 3-4) as the most comprehensive early attempt. Since then, information on the evolution of these bryozoans has increased considerably, and the incorporation of the Austral free-living cluster into the global biogeography of the free-living bryozoans is here briefly summarized in five time slots (Fig. 13), highlighting the unique status of this intriguing group as presented briefly above. The pattern of shifting hotspots for the lunulite bryozoan clades outlined below may, of course, be subject to change in the light of future findings. Notwithstanding, in its present form, it does not seem to conform with the temporal global hotspot pattern presented by Yasuhara et al. (2022).

**Fig. 13.**
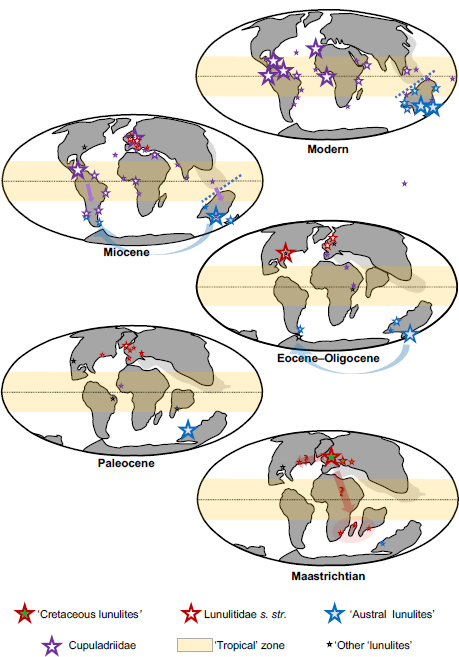
The biogeography of the free-living bryozoans in five easy steps. <u>Maaastrichtian</u>: Total dominance of the Chalk Sea fauna in Northern Europe, thinning rapidly towards south-eastern Europe and Central Asia; questionable connections to taxa in three Gondwana fragments and North America. Note the possible first trace of the Austral free-livings group. <u>Paleocene</u>: The Paleocene ‘dead zone’ marks an over-all low in both density and diversity in the history of the free-living bryozoans, with the Thanetian fauna from NW Western Australia as the outstanding exception. Note the possible first trace of the Cupuladriidae in western Africa. <u>Eocene-Oligocene</u>: Marked re-emergence of lunulites in both North America and Europe, developing a dual, Lunulitidae hotspot, which appear to be separate in both time and space (See Fig. 14). The separate, long-lived Austral hotspot expands to include New Zealand, with early colonizers in southern South America, coexisting with a few representatives of the Cupuladriidae, which are otherwise still only sparingly present in the tropics of Africa and the Mediterranean region. <u>Miocene</u>: The diversity patterns shift dramatically, with a hotspot peak in the Austral realm (now including the southern part of South America), while the Cupuladriidae is rapidly replacing the ‘*Lunulites’* in hot- to warm-water areas across most of the globe. <u>Modern</u>: The Austral free-livings hot-spot reaches its maximum diversity, but they are no longer present in South America. The Cupuladriidae have completely replaced the lunulites across the globe, but still only with a single species in Australia. [Continent positions roughly traced from C.R. Scotese, PALEOMAP Project 2014].

### Maastrichtian

On a global scale, the Chalk Sea fauna in Northern Europe reigns supreme in terms of free-living bryozoan diversity (Håkansson & Voigt 1995 & EH data). At least 100 free-living taxa are known, including a substantial number of specialized forms without setal support (Håkansson 1975). From its northern European hotspot (Fig. 14), this faunal element gradually thins into the south-eastern extent of the Chalk Sea towards Central Asia, where only a few lunulite species are found (Voigt 1967; Favorskaya 1987; Koromyslova & Pervushov 2022). More isolated, undescribed species occurrences are known from North America (P. Taylor pers. comm.) and the three separate Gondwana fragments India (*Lunulites annulata* Stoliczka 1872; Guha & Nathan 1996), Madagascar (*Lunulites pyripora* Canu 1922; Buge 1951), and South Africa (Lunulites sp., Taylor 2019). All three entities had separated from Australia well before the first appearance of any lunulite, with Africa and Madagascar remaining fairly close (Fig. 5). Nevertheless, the three Gondwana species are quite distinct, and none appears obviously related to taxa in the European Chalk Sea. Note also the possible first show of the Austral free-living bryozoans in Northwestern Western Australia referred to briefly above.

**Fig. 14.**
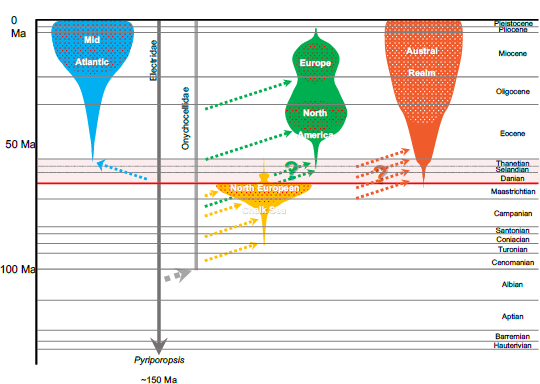
Lunulite hot-spots in time and space, main lunulite clusters color-coded, with hot-spots dotted. The first, and very prominent lunulite hot-spot developed in the North European Chalk Sea (yellow): Highly concentrated, ‘within-basin’, high-frequency iterative evolution, mainly - but not exclusively - from onychocellid roots. Peaks in the Maastrichtian, and terminated in the Maastrichtian-Danian faunal revolution - possibly without descendants. The Lunulitidae s.str.(green) has a longer and more complex story, apparently developing two distinct hots-spots. Firstly, with a North American Gulf & Coastal Plains emphasis, with high densities combined with moderate diversity, possibly iterative evolution from onychocellid roots; peaks in the Middle to Late Eocene and fade away in the Oligocene. Secondly, a less prominent hotspot develops in Europe in the Miocene, seemingly independent from the North American hotspot. The strictly monophyletic Cupuladriidae (blue) develop a high-diversity, still thriving hotspot centered around the Mid Atlantic early in the Miocene. The origin of the polyphyletic Austral Realm lunulites (red) remains enigmatic due to a long history in plate-tectonic isolation (see Fig. 4); a prominent, still thriving hot-spot was established by the Late Eocene with only a single non-Austral lunulite species present from the Miocene. The earliest cheilostome family (dark grey) and the likely family root of many lunulites (light grey) included. Arrows indicate origin; stipples indicate probable iterative evolution

As the Chalk Sea clades with some probability suffered near total extinction as a result of the two-step collapse of the long-lived northern European carbonate province and its unique chalk environment (Fig. 3), further exacerbated by the global effects of the K-Pg extinction event (Surlyk, 1997; Håkansson & Thomsen, 1999, O’Dea et al., 2011), the founding role of the few non-European Cretaceous clades is far from obvious due to the Paleocene ‘dead zone’.

Why was this fauna limited to the European Chalk Sea when there were many coeval flooded continents elsewhere? The fact that most of these other regions received greater siliciclastic input than in the European seas, obviously plays a role, but the long-lasting stability of the European Chalk Sea environment (30-35 Ma; Surlyk 1997) strongly suggests that the Chalk Sea fauna was uniquely adapted to the chalk habitat itself (e.g., Surlyk 1972; Håkansson 1975, Håkansson & Thomsen 1999, Heinberg 2007). This is nevertheless surprising in light of the subsequent radiations and dominations of similar siliciclastic shelf habitats by free-living bryozoans in subsequent epochs, and thus requires further exploration.

### Paleocene

The Paleocene ‘dead zone’ marks an over-all global low in the history of the free-living bryozoans (cf. Fig. 3) mirroring well-established declines in diversity in all bryozoan groups (e.g., Håkansson & Thomsen, 1979, 1999; McKinney & Taylor, 2001; Stilwell & Håkansson, 2012; Moharrek et al., 2022) and other marine groups. In Europe, the last Danian remnants of the Chalk Sea province were home to only a few, rare species (Berthelsen 1962; Kvachko 1995), while Selandian and Thanetian free-living bryozoans are even more scarce, with four Thanetian taxa reported from Deep Sea Drilling Project site 117 at Rockall Bank (Cheetham & Håkansson, 1972) constituting a post-Danian Paleocene ‘hotspot’. However, in marked contrast, the Thanetian fauna from Northwestern Western Australia briefly presented above stands out as the single, remarkable exception in terms of both density and diversity. Outside the Austral hotspot there are thus good reasons to explore the notion that no post-Paleocene free-livings have pre-Paleocene ancestors and, therefore, whether the groups ‘Chalk Sea lunulites‘ and Lunulitidae *s. str*. might have coexisted through this comparatively brief interval of time. However, phylogenetic relationships between faunas have not been formally tested.

Somewhat later, in the Paleocene-Eocene, the first rare representatives of the family Cupuladriidae appear in West Africa (Gorodiski & Balavoine 1961) heralding the global Neogene take-over by this group.

### Eocene-Oligocene

Marked re-emergence of typical lunulite bryozoans in around the central North Atlantic rapidly forming a hotspot (Fig. 14), now mostly belonging to the more advanced and more homogenous group Lunulitidae *s. str*. characterized by their porous, aragonitic basal-wall. The co-eval Austral hotspot change to the south-eastern part of the continent towards the end of the Eocene and expands to include first New Zealand and, subsequently, the first, early colonizers in southern South America appear. Early members of the Cupuladriidae were spreading sporadically in Africa and the Mediterranean region, while commencing a strong southward migration in South America coinciding with the Austral invasion.

### Miocene

The diversity and distribution patterns shifted dramatically, with the Cupuladriidae rapidly replacing Lunulitidae s. str. forming a hotspot (Fig. 14) in tropical and sub-tropical waters across most of the globe bar the Austral realm. There, the Austral free-living hotspot reached its peak, comprising different groups which temporarily reached southern part of South America. Only a single cupuladriid species (*Cupuladria guineensis* (Busk, 1854; our Fig. 1E) appears to have successfully crossed the Wallace line into Australia.

### Modern

The two Miocene hotspots persist into the modern faunas, but no free-living bryozoans are any longer present in southern South America. Rapid Late Neogene cooling may have caused their local extinction, but that does not explain why the Austral part of the fauna were not successful in progressing further north in South America in tune with their preferred climate condition.

As we have attempted to illustrate here, the free-living lunulite bryozoans have a complex history of rise and demise of several unrelated groups across many continents since the Late Turonian., They are generally well preserved as fossils, their colonies and zooidal characters assist in high-level taxonomy, they are diverse but not excessively so, have modern representative and are often abundant in marine assemblages. As such, the waxing and waning of the group offers a unique model system to explore evolutionary patterns, and the drivers of radiations. This is especially so given that, as we have shown, the multiple adoption of the free-living mode of life required comparatively few and simple modifications to the colonial life mode of cheilostomes. To explore the drivers of these evolutionary originations and radiations, the next step will be to construct phylogenetic relationships of a few major groups to other bryozoan faunas to document the frequency of convergent evolution. In addition, increased collections focused on shelf faunas where free-living bryozoan are typically found, especially in tropical regions such as the Indo Pacific, Indian Oceana and Tropical Eastern Pacific, would likely greatly enhance the biogeographic structure we have outlined here. It is essential that such collections are accompanied by full taxonomical treatments.

## Acknowledgments

David Haig (The University of Western Australia) read an early version of the manuscript providing useful comments and suggestions. Phil Bock (Melbourne) generously provided most of the images of SE Australian free-living bryozoans, with additional images provided by Mary Spencer-Jones (Natural History Museum, London). A number of SEM images of NW Australian free-livings were kindly facilitated by Helen Ryan (Western Australian Museum).

## REFERENCES

Barber P H, Palumbi S R, Erdmann M V & Moosa M K 2000. Biogeography: a marine Wallace’s line? Nature 406, 692–693.

Beu A G, Griffin M & Maxwell P A 1997. Opening of Drake Passage gateway and Late Miocene to Pleistocene cooling reflected in Southern Ocean molluscan dispersal: evidence from New Zealand and Argentina. Tectonophysics 281, 83–97.

Bock, P. E. 2022 The Bryozoa home page. http://www.bryozoa.net

Bock P E & Cook P L 1998. Otionellidae, a new family including five genera of free-living lunulitiform Bryozoa (Cheilostomatida). Memorie di Scienze Geologiche 50, 195–211.

Bock P E. & Cook P L 1999. Notes on Tertiary and Recent ‘Lunulite’ Bryozoa from Australia. Memorie di Scienze Geologiche, 51, 415–430.

Bone Y & James N P 1993. Bryozoans as carbonate sediment producers on the cool-water Lacepede Shelf, southern Australia. Sedimentary Geology 86, 247–271.

Buge E 1951. Bryozoaires. In: Collignon, M. (Ed.): Faune maestrichtienne de la Côte d’Ambatry (Province de Betioky), Madagascar. Ann. Géeol. Serv. Mines (Gouvernement Général de Madagascar et Dépendantes) 19, 87–88.

Buge E & Muniz G da C B 1974. *Lunulites (Heteractis) barbosae* – nouvelle espèce de bryozoaire lunulitiforme (Bryozoa, Cheilostomata) du Paléocène du nord-est du Brésil. Annales de Paléontologie, Invertébrès, Paris 60, 191–201.

Busk G 1852a. An account of the Polyzoa and sertularian zoophytes collected in the voyage of the "Rattlesnake" on the coast of Australia and the Louisiade Archipelago. In J. Macgillivray, Narrative of the voyage of H.M.S. Rattlesnake commanded by the late Captain O. Stanley during the years 1846-1850.

Busk G 1852b. Catalogue of marine Polyzoa in the collection of the British Museum. I. Cheilostomata. Trustees of the British Museum (Natural History), London. 1–54.

Busk G 1854. Catalogue of marine Polyzoa in the collection of the British Museum, II. Cheilostomata. Trustees of the British Museum (Natural History), London. 55–120.

Busk G 1884. Report on the Polyzoa collected by H.M.S. Challenger during the years 1873–1876. Part 1. The Cheilostomata. Report on the Scientific Results of the Voyage of the H.M.S. “Challenger”, Zoology 10, 1-216.

Canu, F., 1904, Les bryozoaires du Patagonien. Échelle des bryozoaires pour les terrains tertiaires. Mémoires de la Société Géologique de France: Paléontologie, 12, 1–30.

Canu, F., 1908, Iconographie des bryozoaires fossiles de ĺArgentine. Première partie. Anales del Museo Nacional de Buenos Aires, serie 3, 10, 245–341.

Canu F 1922. Bryozoaires. In: Cottreau. Paleontologie de Madagascar. Annales de Paleontologie. Paris, 15-139.

Canu F & Bassler R S 1917. A synopsis of American Early Tertiary Cheilostome Bryozoa. United States National Museum Bulletin 96, 1-87.

Canu F & Bassler R S 1919. Fossil Bryozoa from the West Indies. Publications of the Carnegie Institution 291, 75–102.

Canu F & Bassler R S 1923. North American later Tertiary and Quaternary Bryozoa. United States National Museum Bulletin 125, 1–302.

Casadio S, Nelson C, Taylor P, Griffin M & Gordon D 2010. West Antarctic Rift system: a possible New Zealand-Patagonia paleobiogeographic link. Ameghiniana 47, 129–132.

Chapman F 1913. Descriptions of new and rare fossils obtained by deep boring in the Mallee. Proceedings of the Royal Society of Victoria 26, 165–191.

Cheetham A H & Håkansson E 1972. Preliminary report on Bryozoa (Site 117). In: Laughton A S, Berggren W A et al.: Initial Reports of the Deep Sea Drilling Project XII, 432-441 & 476-507.

Cook P L 1965. Notes on some polyzoa with conical zoaria. Cahiers de Biologie Marine 6, 435–454.

Cook P L & Chimonides P J 1978. Observations on living colonies of *Selenaria* (Bryozoa, Cheilostomata) I. Cahiers de Biologie Marine 19,147–158.

Cook P L & Chimonides P J 1983. A short history of the lunulite bryozoa. Bulletin of Marine Science 33(3), 566–581.

Cook P L & Chimonides P J 1984a. Recent and fossil Lunulitidae (Bryozoa, Cheilostomata), 1. The genus *Otionella* from New Zealand. Journal of Natural History 18, 227-254.

Cook P L & Chimonides P J 1984b. Recent and fossil Lunulitidae (Bryozoa, Cheilostomata), 2. Species of *Helixotionella* gen. nov. from Australia. Journal of Natural History 18, 255-270.

Cook P L & Chimonides P J 1985a. Recent and fossil Lunulitidae (Bryozoa, Cheilostomata), 3. ‘Opesiulate’ and other species of *Selenaria*, sensu lato. Journal of Natural History 19, 285-322.

Cook P L & Chimonides P J 1985b. Recent and fossil Lunulitidae (Bryozoa, Cheilostomata), 5. *Selenaria alata* Tenison Woods and related species. Journal of Natural History 19, 337-358.

Cook P L & Chimonides P J 1985c. Recent and fossil Lunulitidae (Bryozoa, Cheilostomata), 4. American and Australian species of *Otionella*. Journal of Natural History 19, 575-603.

Cook P L & Chimonides P J 1986. Recent and fossil Lunulitidae (Bryozoa, Cheilostomata), 6. *Lunulites* sensu lato and the genus *Lunularia* from Australasia. Journal of Natural History 20, 681–705.

Cook P L & Chimonides P J 1987. Recent and fossil Lunulitidae (Bryozoa, Cheilostomata), 7. *Selenaria maculata* (Busk) and allied species from Australasia. Journal of natural history 21, 933–966.

Cook P L & Chimonides P J 1994. Notes on the family Cupuladriidae (Bryozoa), and on *Cupuladria remota* sp.n. from the Marquesas Islands. Zoologica Scripta 23/3, 251–268.

Cook P L, Bock P E, Gordon D P & Weaver H J (editors) 2018a: Australian Bryozoa Volume 1. Biology, Ecology and Natural History. SIRO Publishing, Melbourne, 200 pp.

Cook P L, Bock P E, Gordon D P & Weaver H J (editors) 2018b: Australian Bryozoa Volume 2, Taxonomy of Australian Families. CSIRO Publishing, Melbourne, 320 pp.

de Blainville H M D 1830. Zoophytes. Pages 535-546 in G F Cuvier editor Dictionnaire des Sciences naturelles, dans lequel on trate méthodiquement des differents êtres de la nature … par plusieur professeurs du Muséum Nationale d’Histoire Naturelle et des autres principales écoles de Paris 60. F G Levrault, Paris.

Dalziel I W D, Lawver L A, Pearce J A, Barker P F, Hastie A R, Barfod D N, Schenke H-W & Davis M B 2013. A potential barrier to deep Antarctic circumpolar flow until the late Miocene? Geology 41(9), 947–950.

Dick M H, Herrera Cubilla A & Jackson J B C 2003. Molecular phylogeny and phylogeography of free-living Bryozoa (Cupuladriidae) from both sides of the Isthmus of Panama Molecular Phylogenetics and Evolution 27, 355–371.

Di Martino E, Taylor P D, Cotton L J & Pearson P N 2017. First bryozoan fauna from the Eocene–Oligocene transition in Tanzania. Journal of Systematic Palaeontology, https://doi.org/10.1080/14772019.2017.1284163

Di Martino E, Greene S & Taylor P D 2018. Discoradius, a new name for the genus Heteractis Gabb & Horn, 1862 (Cheilostomata, Bryozoa), junior homonym of *Heteractis* Milne-Edwards & Haime, 1851 (Cnidaria, Actiniaria). Journal of Systematic Palaeontology 16 (5), p 445.

Driscoll E G, Gibson J W & Mitchell S W 1971. Larval selection of substrate by the Bryozoa *Discoporella* and *Cupuladria*. Hydrobiologia, 37, 347–359.

Etheridge R Jr 1901. Additional notes on the palaeontology of Queensland (Part 2). Bulletin of the Geological Survey of Queensland 13, 1–37.

Favorskaya T A 1987. Bryozoans of the Maastrichtian of Eastern Turkmenistan. Yearbook All-Union Paleontological Society XXX, 82-107.

Gordon D P, Bock P E, Souto-Derungs J & Reverter-Gil O 2019. A bryozoan tale of two continents: faunistic data for the Recent Bryozoa of Greater Australia (Sahul) and Zealandia, with European comparisons. Austraslasian Palaeontological Memoirs 52, 13–22.

Gorodiski A & Balavoine P 1961. Bryozoaires crétacés et éocènes du Sénégal. Bulletin du Bureau de Recherches Géologiques et Minières 4, 1–15.

Greeley R 1967. Natural orientation of lunulitiform bryozoans. Bulletin of the Geological Society of America 78, 1179–1182.

Guha A K & Nathan D S 1996. Bryozoan fauna of the Ariyalur Group (Late Cretaceous), Tamilnadu and Pondicherry, India. Palaeontologia Indica, new series 49, 215 pp.

Hagenow von F 1839. Monographie der Rügen’schen Kreide-Versteinerungen. Abt. 1. Phytolithen und Polyparien. Neues Jahrbuch für Mineralogie 1839, 252–296.

Haig D W, Smith M & Apthorpe M C 1997. Middle Eocene Foraminifera from the type Giralia Calcarenite, Gascoyne Platform, Southern Carnarvon Basin, Western Australia. Alcheringa 21, 229–245.

Håkansson E 1975. Population structure of colonial organisms. A palaeoecological study of some free-living Cretaceous bryozoans. Documents des Laboratoires de Géologie de la Faculté Sciences de Lyon H.S. 3(2), 385–399.

Håkansson E & Thomsen E 1979. Distribution and types of bryozoan communities at the boundary in Denmark. In Birkelund T & Bromley R G, editors Cretaceous-Tertiary boundary events. Symposium, Copenhagen, September 1979, 1. The Maastrichtian and Danian of Denmark. University of Copenhagen, Copenhagen. 78–91.

Håkansson E & Thomsen E 1999. Benthic extinction and recovery patterns at the K/T boundary in shallow water carbonates. Palaeogeography Palaeoclimatology Palaeoecology 154, 67–85.

Håkansson E & Thomsen E 2001. Asexual propagation in cheilostome Bryozoa: evolutionary trends in a major group of colonial animals. In Jackson J B C, Lidgard S & Mckinney F K (eds). Evolutionary patterns: growth, form and tempo in the fossil record. University of Chicago Press, Chicago, IL,. 326–347.

Håkansson E & Voigt E 1995. New free-living bryozoans from the northwest European Chalk. Bulletin of the Geological Society of Denmark 4, 187–207.

Håkansson E & Winston J E 1985. Interstitial bryozoans: unexpected life forms in a high energy environment. In Nielsen C & Larwood G P editors Bryozoa: Ordovician to Recent. Olsen & Olsen, Fredensborg. 125-134

Håkansson E, Bromley R & Perch-Nielsen K 1974. Maastrichtian chalk of north-west Europe – a pelagic shelf sediment. International Association of Sedimentologists, Special Publications 1, 211–233

Håkansson E, Gordon D P & TAYLOR P D in press. Bryozoa from the Maastrichtian Korojon Formation, Western Australia. Fossils and Strata.

Håkansson E, O’Dea A & Rosso A 2019. The Free-Living Cheilostome Bryozoans – pursuing the unobtainable. 18 International Bryozoology Association Conference, Liberec, 16-22 June 2019.

Hara U, Mörs T, Hagström J & Reguero MA 2018. Eocene bryozoan assemblages from the La Meseta Formation of Seymour Island, Antarctica. Geological Quarterly 62 (3), 705–728.

Heinberg C 2007. Evolutionary ecology of nine sympatric species of the pelecypod *Limopsis* in Cretaceous chalk. Lethaia 12(4), 325–340.

Hincks T 1881a. Contributions towards a general history of the marine Polyzoa. Part VI. Polyzoa from Bass’s Straits. Annals & Magazine of Natural History (Series 5) 8,1-14 & 122–129.

Hincks T 1881b. On a collection of Polyzoa, from Bass’s Straits, presented by Capt. Cawne Warren to the Liverpool Free Museum. Proceedings of the Literary and Philosophical Society of Liverpool 35, 249–270.

Hincks T 1882. Contributions towards a general history of the marine Polyzoa. Part X. Foreign Cheilostomata (miscellaneous). Annals & Magazine of Natural History (Series 5) 10,160–170.

Hodel F., Grespan R, de Rafélis M, Dera G, Lezin C, Nardin E, Rouby D, Aretz M, Steinnman M, Buatier M, Lacan F, Jeandel C & Chavagnac V 2021. Drake Passage gateway opening and Antarctic Circumpolar Current onset 31 Ma ago: The message of foraminifera and reconsideration of the Neodymium isotope record. Chemical Geology 570, 120171, 1–14.

Koromyslova A V & Pervushov E 2022. Uppermost Turonian bryozoans from the Lower Volga River region: scanning electron microscopy and micro-computed tomography studies. Neues Jahrbuch für Geologie und Paläontologie – Abhandlungen 305(3), 263–95.

Kvachko V I 1995. Late Cretaceous and Paleocene bryozoans of the genus *Lunulites* from the Middle Volga, Crimea and Mangyshlak. Paleontological Journal 29(4), 36–45

Lagaaij R 1952. The Pliocene Bryozoa of the Low Countries and their bearing on the marine stratigraphy of the North Sea region. Mededelingen van de Geologische Stichting 5, 233pp.

Lagaiij R 1963. *Cupuladria canariensis* – portrait of a bryozoan. Palaeontology 6, 172–217.

Lamarck J B P A de M de 1816. Histoire naturelle des Animaux sans Vertèbres … précédée *d’une introduction offrant la détermination des caractéres essentiels de l’animal, sa* distinction du végétal et des autres corps naturels, enfin, exposition des principes fondamentaux de la zoologie. Verdiere, Pari. 568pp.

Lawver L A & Gahagan L M 2003. Evolution of Cenozoic seaways in the circum-Antarctic region. Palaeogeography, Palaeoclimatology, Palaeoecology 198, 11–37.

Levinsen G M R 1909. Morphological and systematic studies on the cheilostomatous Bryozoa, Nationale Forfatterers Forlag, Copenhagen. 431pp.

López-Gappa J, Pérez L M & Griffin M 2017. First record of a fossil selenariid bryozoan in South America. Alcheringa, 41 (3): 365–368.

MacGillivray P H 1881. Polyzoa. In Prodromus of the Zoology of Victoria (Ed. F McCoy) 1(6), 27-46.

MacGillivray P H 1886. Polyzoa. In Prodromus of the Zoology of Victoria (Ed. F McCoy) 2(12), 63-73.

MacGillivray P H 1895. A monograph of the Tertiary Polyzoa of Victoria. Transactions of the Royal Society of Victoria 4,1–166.

Maplestone C M 1904. Notes on the Victorian fossil Selenariidae, and descriptions of some new species (Recent and fossil). Proceedings of the Royal Society of Victoria (new series) 16, 207–217.

Maplestone C M 1911. The results of deep-sea investigations in the Tasman Sea. 1 – The expedition of the H.M.C.S. "Miner". 5 - Polyzoa supplement. Records of the Australian Museum 8, 118–119.

Marcus E & Marcus E 1962. On some lunulitiform Bryozoa. Boletimda da Faculdade de Filosofia, Ciências e Letras, Universidade de São Paulo, Zoologia 24, 281-342.

Marsson, T. 1887: Die bryozoen der weissen schreibkreideder insel Rügen. Palæontologische Abhandlungen herausgegeben von W. Dames und E. Kayser 4/1, 112 pp.

Maturo F J S 1968. The distributional pattern of the bryozoa of the East coast of the United States exclusive of New England. Atti della Società Italiana di Scienze Naturali e del Museo Civico di Storia Naturale di Milano 108, 261-284.

McKinney F K & Taylor P D 2001. Bryozoan generic extinctions and originations during the last one hundred million years. Palaeontologia Electronica 4 (1), 26pp. http://palaeo-electronica.org/2001_1/bryozoan/issue1_01.htm

Michelotti G 1838. Specimen zoophytologiæ diluvianæ. Turin 227 pp.

Moharrek F, Taylor P D, Silvestro D, Jenkins H L, Gordon D-P & Waeschenbach A 2022. Diversification dynamics of cheilostome bryozoa based on a bayesian analysis of the fossil record. Palaeontology https://doi: 10.1111/pala.12586

Müller R D, Zhirovic S, Williams S E, Cannon J, Seton M, Bower D J, Tetley M G, Heine C, Le Breton E, Liu S, Russel S H J, Yang T, Leonard J & Gurnis M 2019. A Global Plate Model Including Lithospheric Deformation Along Major Rifts and Orogens Since the Triassic. Tectonics 38, 1884–1907

O’Dea A 2009. Relation of form to life habit in free-living cupuladriid bryozoans. Aquatic Biology 7, 1–18.

O’Dea, A., Håkansson, E., Taylor, P. & Okamura, B. 2011: Environmental change prior to the K-T boundary inferred from temporal variation in the morphology of cheilostome bryozoans. Palaeogeography, Palaeoclimatology, Palaeoecology, Palaeoecology, 308, 502–512.

O’Dea A, Jackson J B C, Taylor P D & Rodríguez F 2008. Mode of reproduction in fossil and recent cupuladriid bryozoans. Paleontology 51, 847–864.

Ostrovsky A N 2013. Evolution of Sexual Reproduction in Marine Invertebrates. Example of Gymnolaemate Bryozoans. Ch. 2 Cheilostome Brood Chambers: Structure, Formation, Evolution. Springer, Dordrecht. 115-228. https://link.springer.com/book/10.1007%2F978-94-007-7146-8

Ostrovsky A N, O’Dea A & Rodríguez F 2009. Comparative Anatomy of Internal Incubational Sacs in Cupuladriid Bryozoans and the Evolution of Brooding in Free-Living Cheilostomes. Journal of Morphology 270, 1413–1430.

Parker S A, & Cook P L 1994. Records of the Bryozoan family Selenariidae from Western Australia and South Australia, with the descriptions of new species of *Selenaria*. Records of the South Australian Museum 27, 1–11.

Philippi R A 1887. Die Terrtiaren und Quartaren Versteinerungen Chiles. Leipzig, F A Brockhaus. 266 pp.

Pohowsky R A 1973. A Jurassic Cheilostome from England. In: Larwood G P (ed.) Living and Fossil Bryozoa. Academic Press, London and New York. 447–461.

Scher H D & Martin E E 2006. Timing and Climatic Consequences of the Opening of Drake Passage. Science, 312, 428.

Schmidt R 2007. Australian Cainozoic Bryozoa, 2: Free-living Cheilostomata of the Eocene St Vincent Basin, S.A., including *Bonellina* gen nov. Alcheringa 31, 67–84.

Scotese C R 2014. Atlas of Neogene Paleogeographic Maps (Mollweide Projection), Maps 1-7, Volumes 1, The Cenoaoic, PALEOMAP Atlas for ArcGIS, PALEOMAP Project, Evanston, IL.

Stilwell J D & Håkansson E 2012. Survival, but ! New tales of ‘Dead clades walking’ from Austral and Boreal Post-K-T Assemblages. In: Talent J D (Ed.): Earth and Life, International Year of Planet Earth. Springer Science. 795-810. https://doi: 10.1007/978-90-481-3428-1_26

Stoliczka F 1872. The Cretaceous fauna of southern India. The Ciliopoda. Memoir of the Geological Survey of India. Palaeontologica Indica 4, 33–68

Surlyk, F., 1972. Morphological adaptations and population structures of the Danish chalk brachiopods (Maastrichtian). Det Kongelige Danske Videnskabernes Selskab. Biologiske Skrifter 19, 57pp.

Surlyk F 1997. A cool-water carbonate ramp with bryozoan mounds: Late Cretaceous-Danian of the Danish Basin. In: James N P & Clarke J D A (eds.): Cool-water carbonates. SEPM Special Publication 56, 293-307.

Taylor P D 1988. Major radiation of cheilostome bryozoans: Triggered by the evolution of a new larval type? Historical Biology, 1(1) 45–64.

Taylor P D 2019. A brief review of the scanty recordof Cretaceous bryozoans from Gondwana. Australian Palaeontological Memoir 52, 147–154.

Tenison Woods JE 1865. On some Tertiary fossils in South Australia. Transactions and Proceedings of the Royal Society of Victoria 6, 3-6.

Tenison Woods J E 1880. On some Recent and fossil species of Australian Selenariidae (Polyzoa). Transactions of the Royal Society of South Australia, Adelaide 3, 1–12.

Voigt E 1967. Oberkreide-Bryozoen aus den asiatischen Gebieten der UsSSR. Mitteilungen Geologishen Staatsinstitut Hamburg 36, 5–95.

Winston J E 1988. Life histories of free-living bryozoans. National Geographic Research, 4, 528–539.

Winston J E & Håkansson E 1986. The interstitial bryozoan fauna from Capron Shoal, Florida. American Museum Novitates 2865, 1–50.

Yasuhara M, Huang H H M, Reuter M, Tian S Y, Cybulski J D, O’Dea A, Mamo B L, Cotton L J, Di Martino E, Feng R & Tabor C R 2022. Hotspots of Cenozoic tropical marine biodiversity. In: Hawkins S J et al. (Eds). Oceanography and Marine Biology: An Annual Review, 60. CRC Press. 243–300.

Yang J, Lan T, Zhang X & Martin R. Smith M R 2023. *Protomelission* is an early dasyclad alga and not a Cambrian bryozoan. Nature 615, 468–471. https://doi.org/10.1038/s41586-023-05775-5

Zhang Zhil., Zangh Zhif., Ma J, Taylor P D, Strotz L C, Jacquet S M, Skovsted C B, Chen F, Han J & Brock G A 2021. Fossil evidence unveils an early Cambrian origin for Bryozoa. Nature 599, 251–255. https://doi.org/10.1038/s41586-021-04033-w

